# Observation of the dynamic changes in the urinary proteome in rats during immunization

**DOI:** 10.1101/2022.12.10.519864

**Authors:** Yunlong Wang, Youhe Gao

## Abstract

**Objective:** Changes in the immune system in the urine proteome were observed by injecting bovine serum albumin and aluminum hydroxide adjuvant into rats.

**Methods:** In this study, bovine serum albumin and aluminum hydroxide adjuvant were injected into rat thigh muscle, urine was collected, differential proteins were identified by liquid chromatography—mass spectrometry (LC—MSMS/MS), and biological pathways of differential proteins were analyzed by IPA software to observe the changes in the immune system as evidenced by rat urinary proteins.

**Results:** Fifteen rats were intramuscularly injected with normal saline, aluminum hydroxide adjuvant, bovine serum albumin, aluminum hydroxide adjuvant, and bovine serum albumin (BSA) mixture to construct the models of the control, adjuvant, BSA, and mixed groups. Upon comparing the different proteins between different groups to obtain the relevant biological pathways, it was found that adjuvants can be observed in urine to help bovine serum albumin stimulate the immune system to respond earlier. It was also observed in urine that the mixed group successively stimulated immune-related pathways, such as the inflammatory response, T-cell activation, antigen-presenting cell-related pathways, and B-cell-related pathways.

**Discussion:** We can observe changes in the immune system from the urine proteome in the early stage, providing some new clues and a basis for future research on the immune system and accelerating vaccine research and development.

## 1 Introduction

Because urine is not a body fluid, it is an aggregate of biological information that can be easily overlooked. Since urine is not regulated by homeostatic mechanisms, it can reflect many changes and is a place rich in information about such changes, which is beneficial to our research and discovery of early biomarkers^[1]^. Under normal physiological conditions, human urine proteins are relatively stable. If these stable proteins undergo great changes under certain conditions, these proteins can be regarded as good biomarkers^[2]^. In addition, urine has the advantage of being able to be collected noninvasively and continuously and is produced in large amounts. Therefore, we believe that urine is an ideal sample for identifying and studying biomarkers.

In our existing studies, whether it is the study of tumor models such as astrocytoma^[4]^, pancreatic cancer^[5]^, bladder cancer^[6]^, or the study of different kinds of bacteria through intraperitoneal injection^[7]^, significant differential proteins were found in the urine, and we further observed in the urine that the state of the immune system was different each time and would undergo different changes. As early as 2,000 years ago, the famous Greek physician Hippocrates declared that “the best doctor of mankind is himself”, which pointed out that human medicine should change from external interventional treatment to human autoimmunity research^[8]^. Therefore, in urine, we can look for the most basic aspects of the immune system and explore the protein changes related to the immune system and also explore the significance of these differences. This is a new attempt and exploration in the field of urine proteins. In this study, we used normal saline, aluminum hydroxide adjuvant, bovine serum albumin, and a mixture of bovine serum albumin and aluminum hydroxide adjuvant for intramuscular injection in rats. Urine was collected on Days 5, 7, and 14. The collected urine was subjected to protein extraction, enzyme digestion, and mass spectrometry analysis to observe the changes in histones and biological pathways at different time points, providing clues for studying the immune system based on urine protein profiles and for accelerating the development of vaccines.

## 2 Materials and methods

### 2.1 Experimental animals and model construction

Fifteen 150 g male Wistar rats were purchased from Beijing Weitong Lihua Laboratory Animal Technology Co., Ltd. and were divided into a normal saline group (n=3), an aluminum hydroxide adjuvant group (n=4), a bovine serum albumin group (n=4), and a bovine serum albumin and aluminum hydroxide adjuvant mixed group (n=4). The bovine serum albumin group was injected with a dose of 4 mg, the mixed group was injected with the same amount of bovine serum albumin and aluminum hydroxide adjuvant in a 1:1 solution, the aluminum hydroxide adjuvant group was injected with the same amount of aluminum hydroxide adjuvant, and the normal saline group was injected with the same amount of normal saline. All groups were injected uniformly by intramuscular injection in the right thighs of the rats. During the experimental period, the animals were reared under standard conditions of 12 hours of normal light-dark cycle, temperature (22 °C ± 1 °C), and humidity (65%–70%). Urine was collected once a day, proteins were extracted from the urine collected from the rats on the 1st, 3rd, 5th, 7th, and 14th days after injection, and enzyme digestion was performed. All experimental procedures met the standards of animal ethics review. The animal license was SCXK (Beijing) 2016-0006. All experiments were approved by the Institutional Animal Care, Use and Welfare Committee, Institute of Basic Medicine, Peking Union Medical College (Animal Welfare Assurance Number: ACUC-A02-2014-007)

### 2.2 Urine collection and sample processing

#### (1) Urine collection

Following the intramuscular injection of the rats, urine was collected once a day for the following week, followed by a final urine collection on Day 14. Urine was collected from each rat for 10 hours overnight in a metabolic cage without water and food provided. The collected urine was immediately stored at −80 °C the next morning, and the urine samples on the 1^st^, 3^rd^, 5^th^, 7^th^, and 14^th^ days were finally selected for subsequent experiments.

#### (2) Urine protein extraction and enzyme digestion

Urine protein extraction: Urine was centrifuged at 12000 × g, 40 min, and 4 °C to obtain the supernatant. Then, 500 μl of the supernatant was transferred to a new EP tube, and precooled ethanol was added according to the ratio of supernatant:ethanol=1:3 and was stirred evenly, after which it was stored overnight at −20 °C for 12 h. The next day, the solution was mixed, centrifuged at 12000 × g, 30 min, 4 °C to discard the supernatant. The precipitate was inverted onto filter paper and dried with a blower in cold air. Lysate was added (37.5 μl), and the mixture was blown evenly with a pipette tip until there was no precipitation, after which it was centrifuged at 12000 × g, 30 min, 4 °C. The supernatant was then placed a new EP tube and stored at −80 °C. After reconstitution, the protein concentration was determined by the Bradford method.

Urine proteolysis: Urine proteolysis was performed using the FASP method^[9]^. One hundred micrograms of urine protein was added to the filter membrane of a 10 kDa ultrafiltration tube (Pall, Port Washington, NY, USA) using UA solution (8 mol/L urea, 0.1 mol/L Tris-HCl, pH 8.5), and 25 mmol/L NH4HCO3 solutions were washed twice. Trypsin (Trypsin Gold, Promega, Fitchburg, WI, USA) was added according to the ratio of trypsin:protein to 1:50 for digestion, and the cells were digested in a water bath at 37 °C overnight. After overnight incubation, the peptides were collected by centrifugation and passed through an HLB solid-phase extraction column (Waters, Milford, MA) for desalting treatment, dried under vacuum, and stored at −80 °C.

#### (3) LC—MS/MS tandem mass spectrometry analysis

The digested samples were reconstituted in 0.1% formic acid water and diluted to 0.5 μg/μL. Each sample was taken to prepare a mixed peptide sample, which was separated using a high-pH reverse-phase peptide separation kit (Thermo Fisher Scientific). The mixed polypeptide sample was added to a chromatographic column and eluted with a solution of increasing acetonitrile concentration gradient, and ten effluents were collected by centrifugation, dried by a vacuum desiccator, and reconstituted with 0.1% formic acid water. Peptides (Biognosis, Inc.) were synthesized using iRT and added to the ten fractions and each sample at a volume ratio of 10:1. Data acquisition of the 10 fractionated components was performed using an EASY-nLC 1200 ultrahigh-performance liquid chromatography-tandem Orbitrap Fusion Lumos high-resolution mass spectrometer. The peptides dissolved in 0.1% formic acid water were loaded on a precolumn (75 μm×2 cm, 3 μm, C18, 100A°), and the eluent was loaded on a reversed-phase analytical column (50 μm×250 mm, 2 μm, C18, 100A°) with an elution gradient of 4%-35% mobile phase B (80% acetonitrile + 0.1% formic acid + 20% water, flow rate 300 nL/min) for 90 min. For fully automated and sensor signal processing, a calibration kit (iRT kit, Biognosys, Switzerland) was used for all samples at a concentration of 1:20 v/v. The 10 components were analyzed in DDA-MS mode, and the parameters were set as follows: the spray voltage was 2.4 kV, the primary resolution of the Orbitrap was 60000, the scanning range was 350-1550 m/z, the secondary scanning range was 200-2000 m/z, the resolution was 30000, the screening window was 2 Da, and the collision energy was 30% HCD. The AGC target was 5e4, and the maximum injection time was 30 ms. The raw files were constructed and analyzed by PD (Proteome Discoverer 2.1, Thermo Fisher Scientific) software.

#### (4) Mass Spectrometry Data Processing

The PD library search results were used to establish the DIA acquisition method, and the window width and number were calculated according to the m/z distribution density. A single peptide sample was subjected to DIA mode to collect mass spectrometry data. Mass spectral data were processed and analyzed using Spectronaut X software. The raw files collected by each sample DIA were imported to search the library. The high-confidence protein standard was defined as a peptide q value<0.01, and the peak areas of all fragment ions of the secondary peptide were used for protein quantification.

#### (5) Statistical analysis

Missing value filling (KNN method)^[10]^ and CV value screening (CV < 0.3)^[11]^ were performed on the identification results of mass spectrometry, and an independent samples t test was used to compare the data between every two groups. To minimize the influence of rat growth and development on urine proteins, we used the comparison method of adjacent time points, that is, comparisons between the 4th week and the 0th week, the 8th week and the 4th week, and the 12th week and the 8th week. For comparing the 16th week and the 12th week and the 18th week and the 16th week, the criteria for screening differential proteins were as follows: fold change FC≥1.5 or FC≤0.67 between the two groups, P<0.05.

#### (6) Differential protein functional annotation

The screened differential proteins were subjected to functional enrichment analysis using the DAVID database (https://david.ncifcrf.gov/)^[12]^ and IPA software (Ingenuity Systems, Mountain View, CA, USA), all using a P<0.05 significance threshold.

## 3 Experimental results

### 3.1 Analysis of Urine Proteome Changes

#### (1) Analysis of Unsupervised Clustering Results

To intuitively observe the differences between urinary protein groups in the four models, we conducted unsupervised clustering observations of the urinary protein groups at each time point. The unsupervised clustering results are shown in Figure 7, where A, B, C, and D represent the aluminum hydroxide adjuvant (adjuvant), bovine serum albumin (antigen), normal saline (control), and bovine serum albumin and aluminum hydroxide adjuvant (antigen with adjuvant) groups, respectively. For example, B-4-1 represents the first day of rat no. 4, and the other numbers are the same. According to the overall unsupervised clustering results of the four groups, we can roughly see that the saline group was closer to the BSA group, while the aluminum hydroxide adjuvant and antigen plus adjuvant groups were grouped together and significantly separated from the saline and BSA groups. These results suggest that adjuvants may play a role in triggering changes in the immune system.

**Figure 1.**
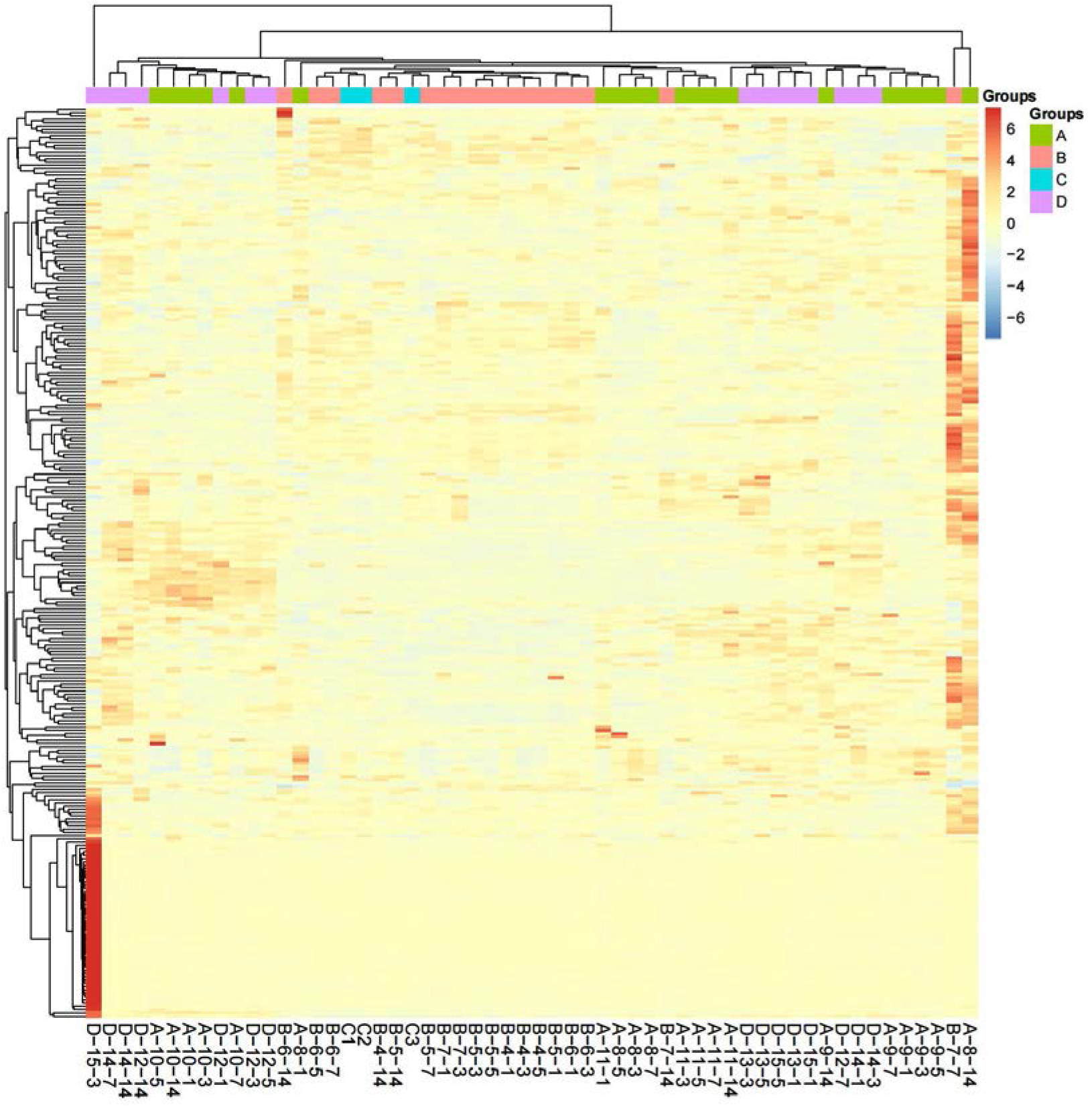
Unsupervised clustering results of urinary protein in 4 groups.

**Figure 2.**
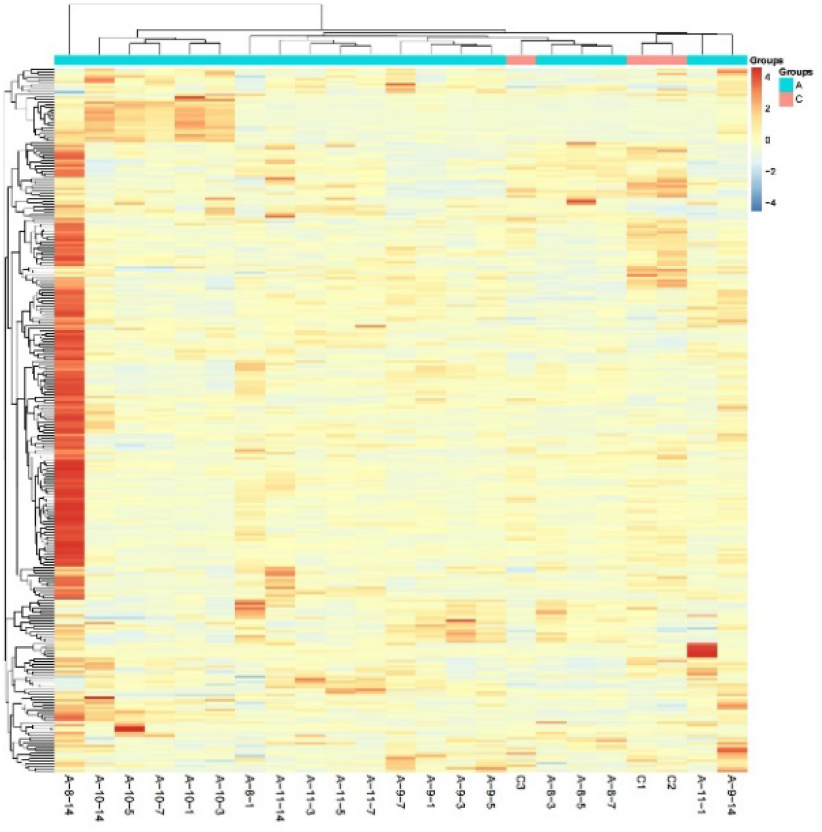
Adjuvant group and control group.

**Figure 3.**
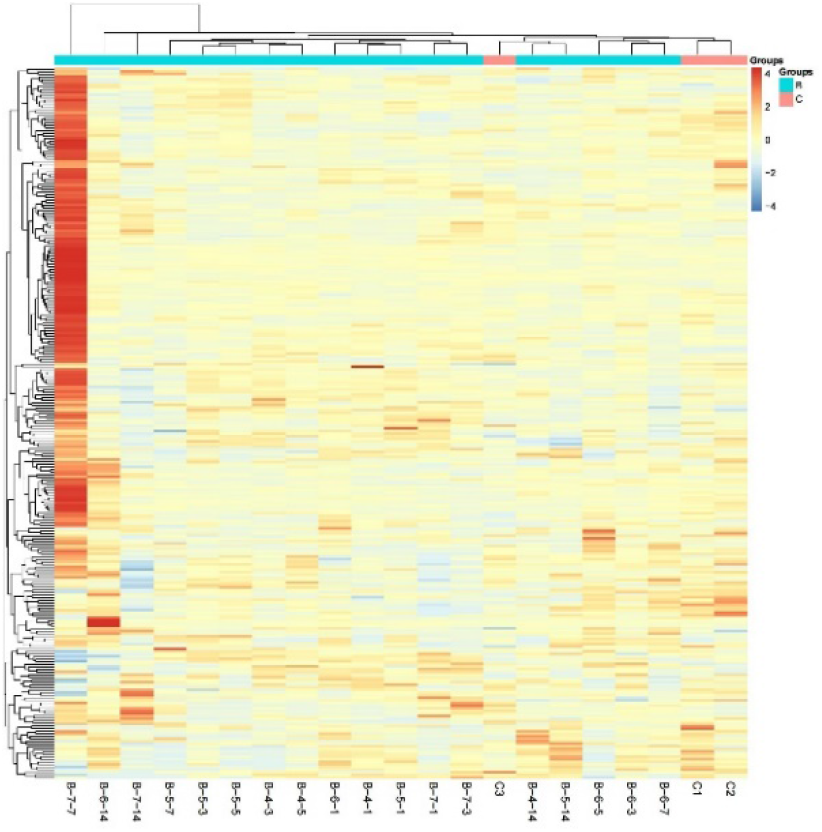
BSA group and control group.

**Figure 4.**
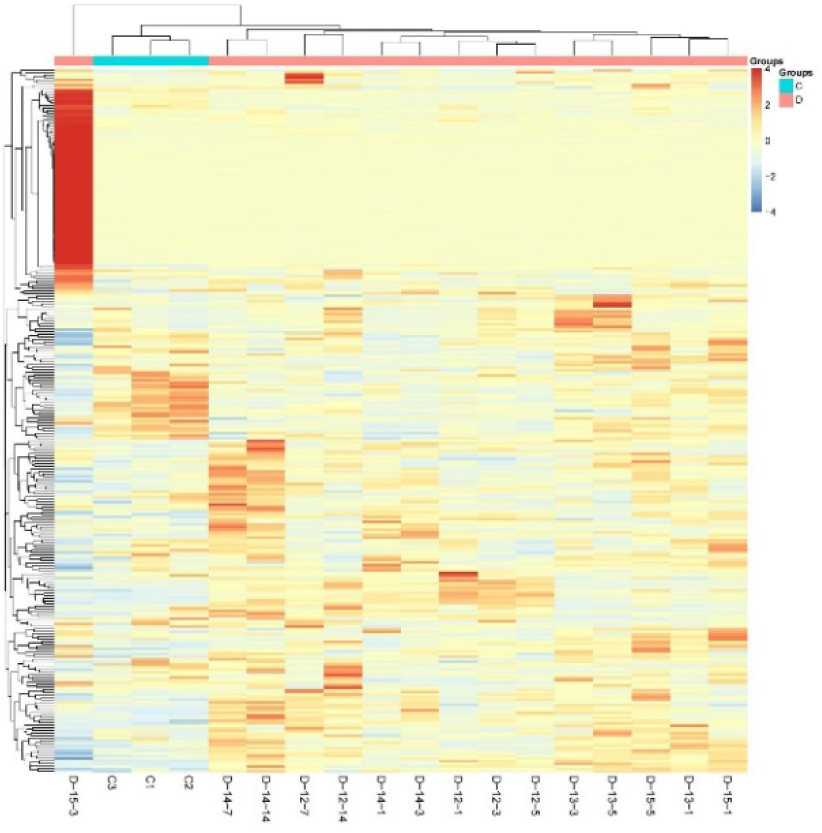
Mixed group and control group.

**Figure 5.**
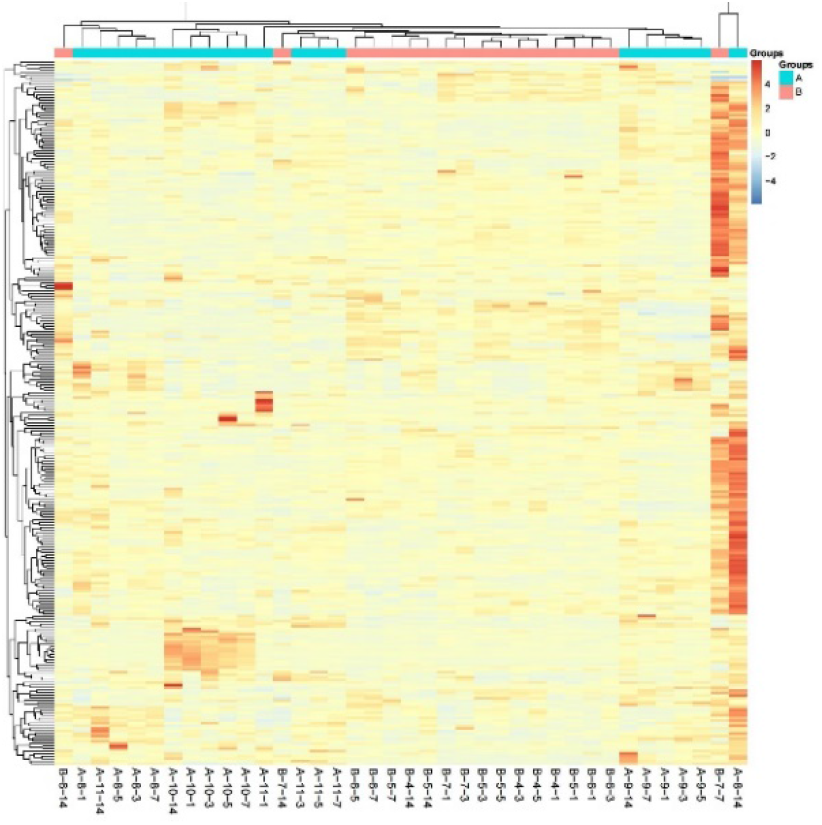
Adjuvant group and BSA group.

**Figure 6.**
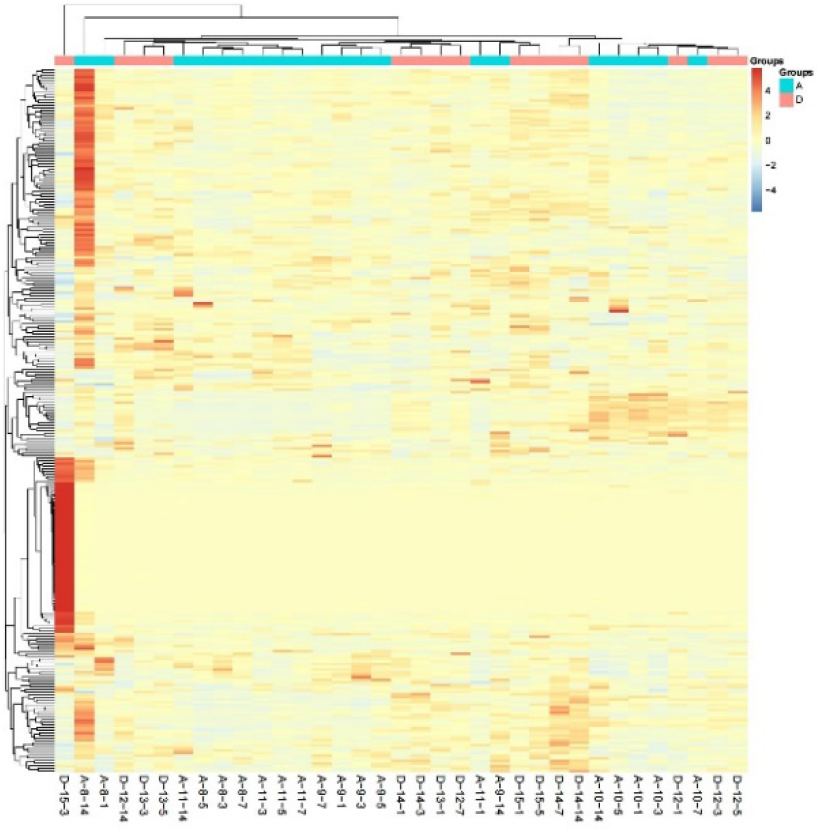
Adjuvant group and mixed group.

**Figure 7.**
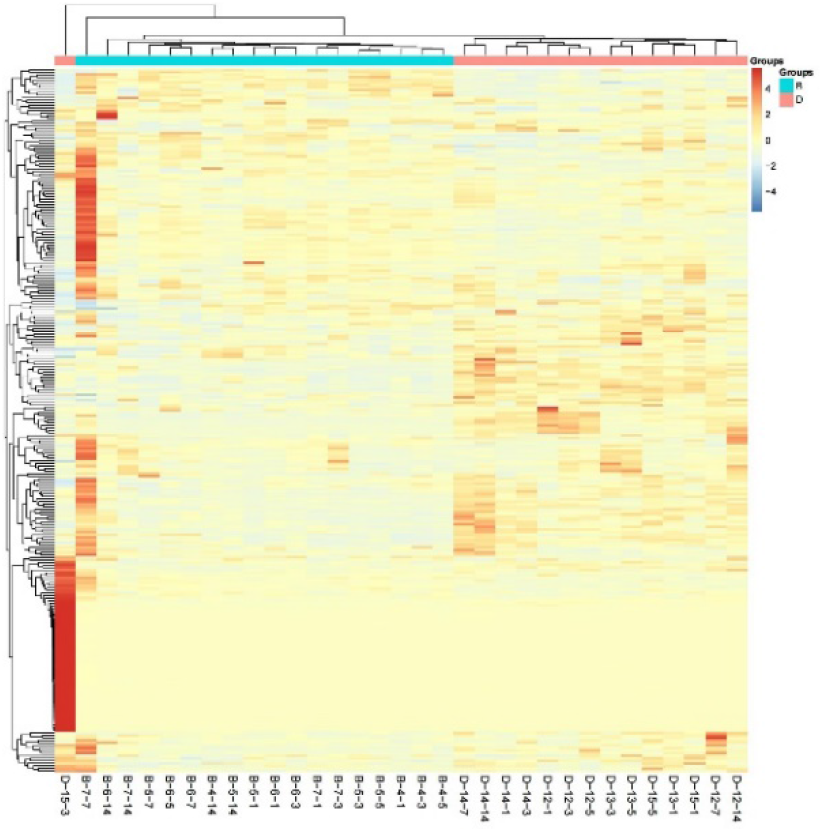
BSA group and mixed group.

Based on the unsupervised clustering of urinary proteins in the 4 groups, we continued to conduct unsupervised clustering by pairwise comparison between the two groups to better observe the overall difference in urinary proteins in each group. The results are shown in Figures 7-12, and the numbers are the same as above. As shown in Figures 8-11, the comparison between our control group and the other three groups shows that the overall trend of the control group was different from each experimental group, but the differences may not be intuitive due to the small number of samples in the control group. As we mentioned before, the unsupervised clustering results of urinary protein in the 4 groups as a whole showed that the addition of adjuvant had a greater effect, and the pairwise comparison results were consistent. As shown in Figures 11 and 13, both the adjuvant group of pure aluminum hydroxide adjuvant and the antigen-adjuvant group of aluminum hydroxide adjuvant mixed with BSA could be clearly divided into two groups by unsupervised clustering. However, as shown in Figure 12, when the adjuvant group was compared with the antigen plus adjuvant group, they were mutually grouped together, and the clustering result was not obvious.

#### (2) Differential protein and biological pathway analyses

We compared the differential proteins of the adjuvant group and the control group, the antigen group, and the control group, the antigen plus adjuvant group, and the antigen plus adjuvant group and the control group, as shown in Table 1. The differentially expressed proteins were further analyzed using Ingenuity Pathway Analysis software, and the IPA pathways compared among different groups at different time points were sorted, as shown in Table 2.

**Table 1.**
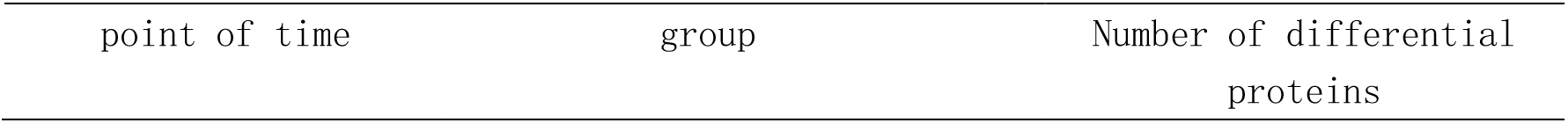
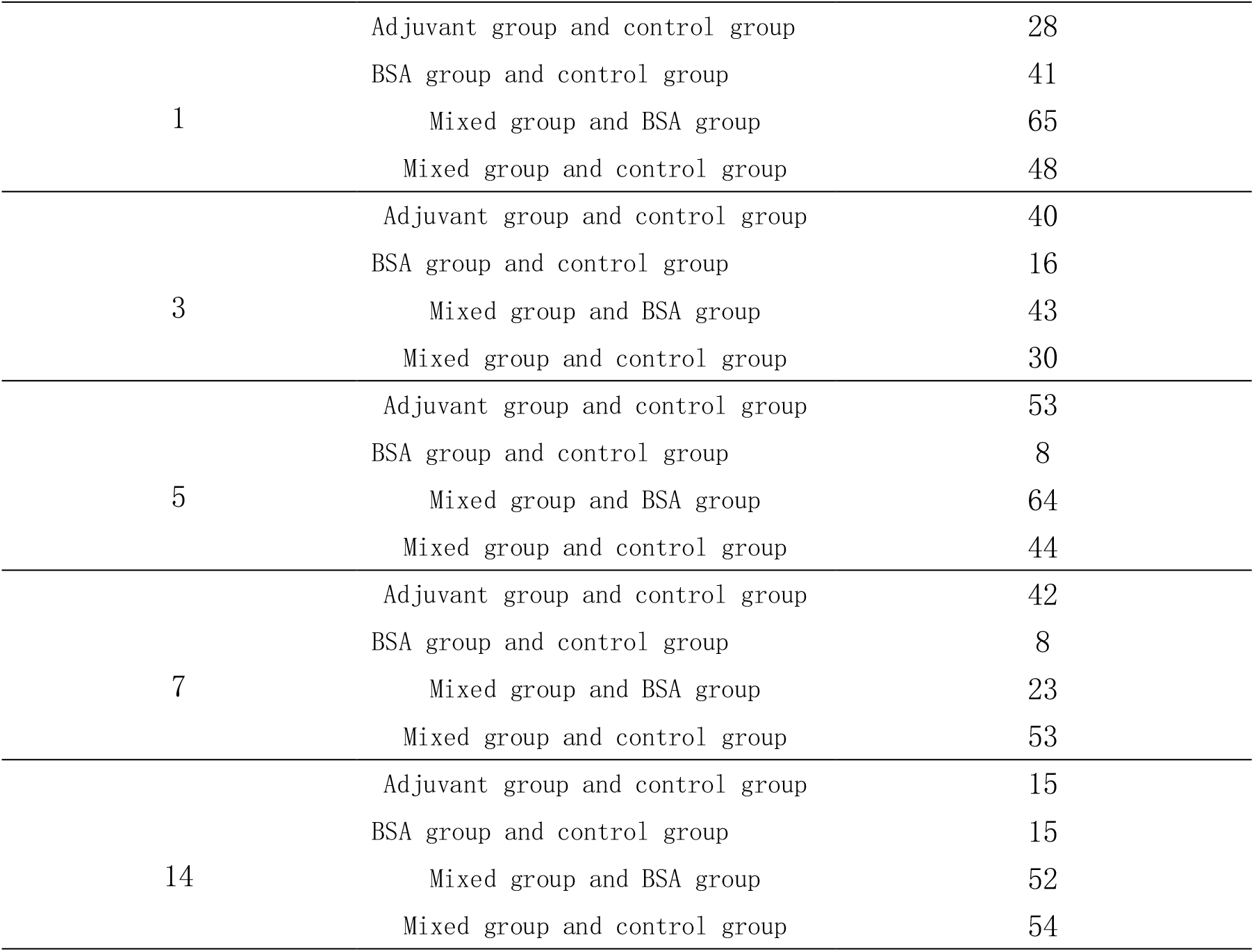
Different proteins in different groups at different time points

| point of time | group | Number of differential proteins |
| --- | --- | --- |
| 1 | Adjuvant group and control group | 28 |
|  | BSA group and control group | 41 |
|  | Mixed group and BSA group | 65 |
|  | Mixed group and control group | 48 |
| 3 | Adjuvant group and control group | 40 |
|  | BSA group and control group | 16 |
|  | Mixed group and BSA group | 43 |
|  | Mixed group and control group | 30 |
| 5 | Adjuvant group and control group | 53 |
|  | BSA group and control group | 8 |
|  | Mixed group and BSA group | 64 |
|  | Mixed group and control group | 44 |
| 7 | Adjuvant group and control group | 42 |
|  | BSA group and control group | 8 |
|  | Mixed group and BSA group | 23 |
|  | Mixed group and control group | 53 |
| 14 | Adjuvant group and control group | 15 |
|  | BSA group and control group | 15 |
|  | Mixed group and BSA group | 52 |
|  | Mixed group and control group | 54 |

**Table 2.**
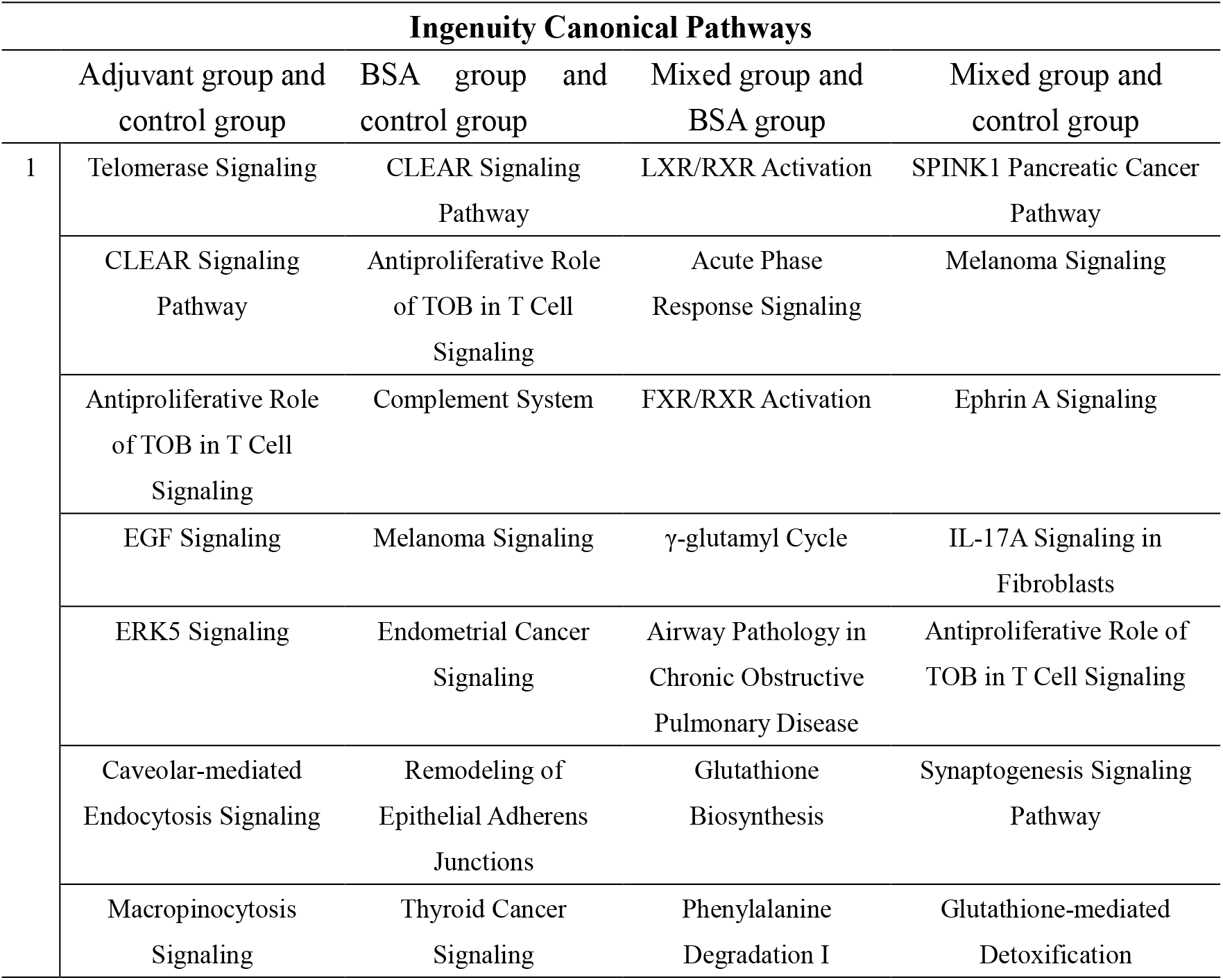
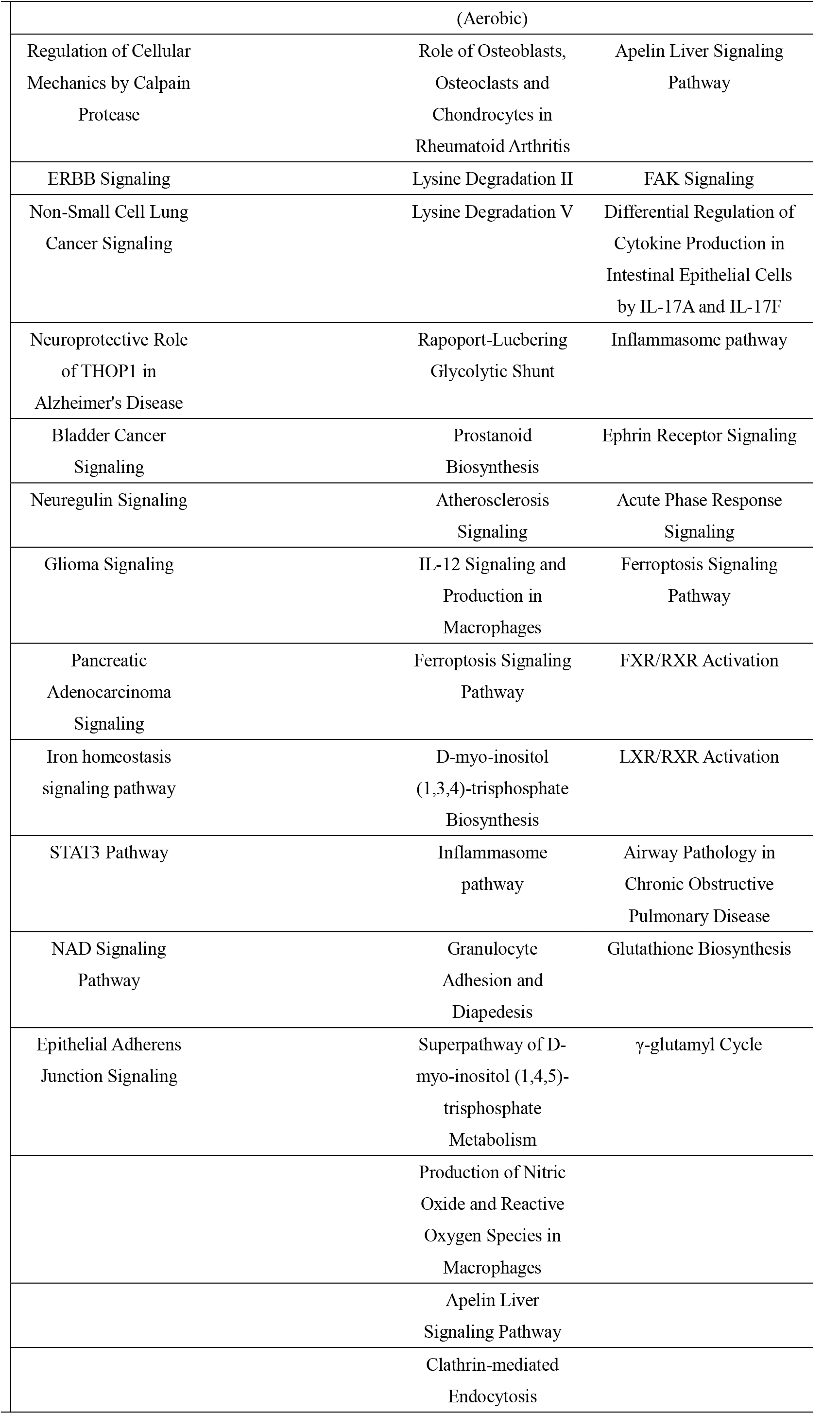
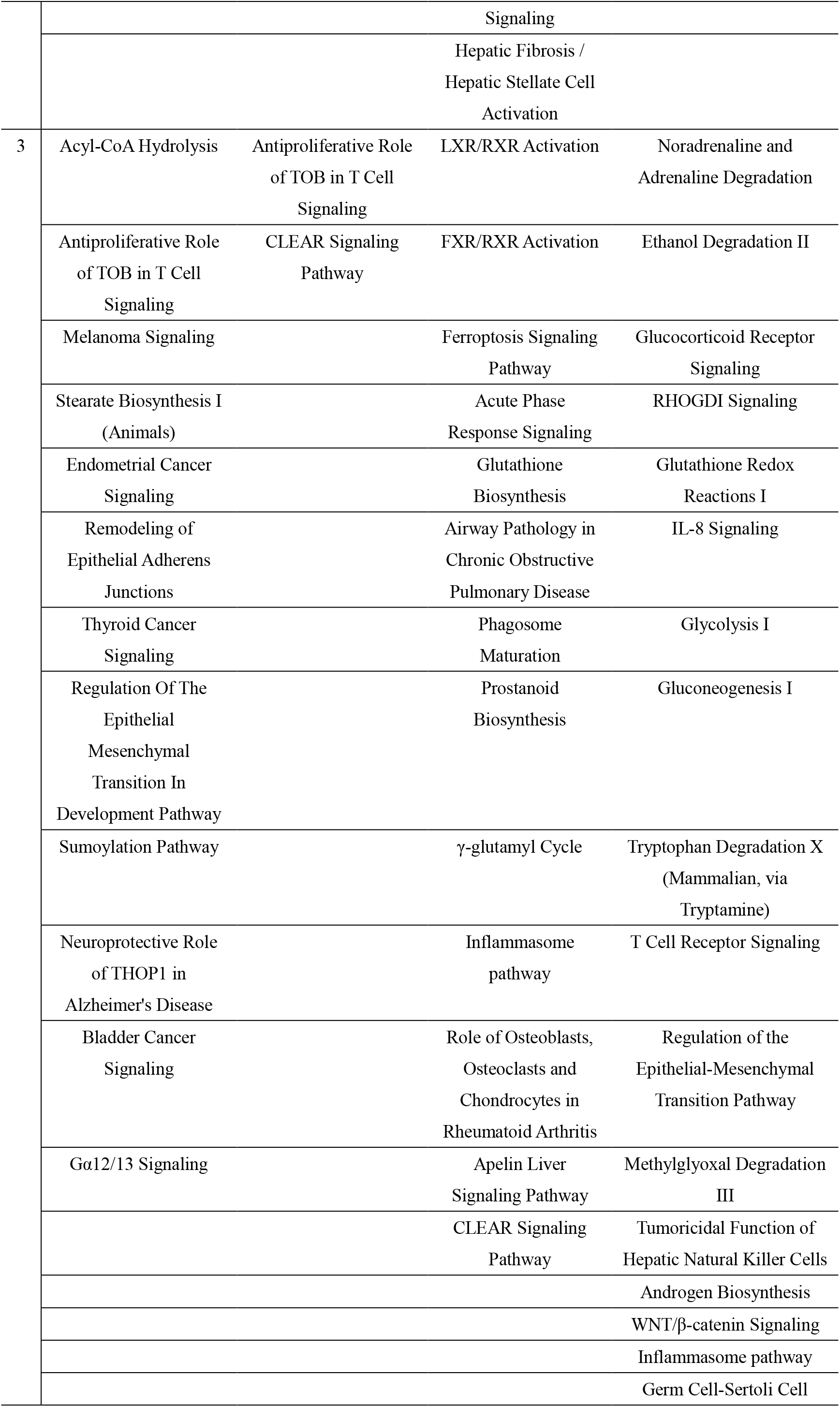
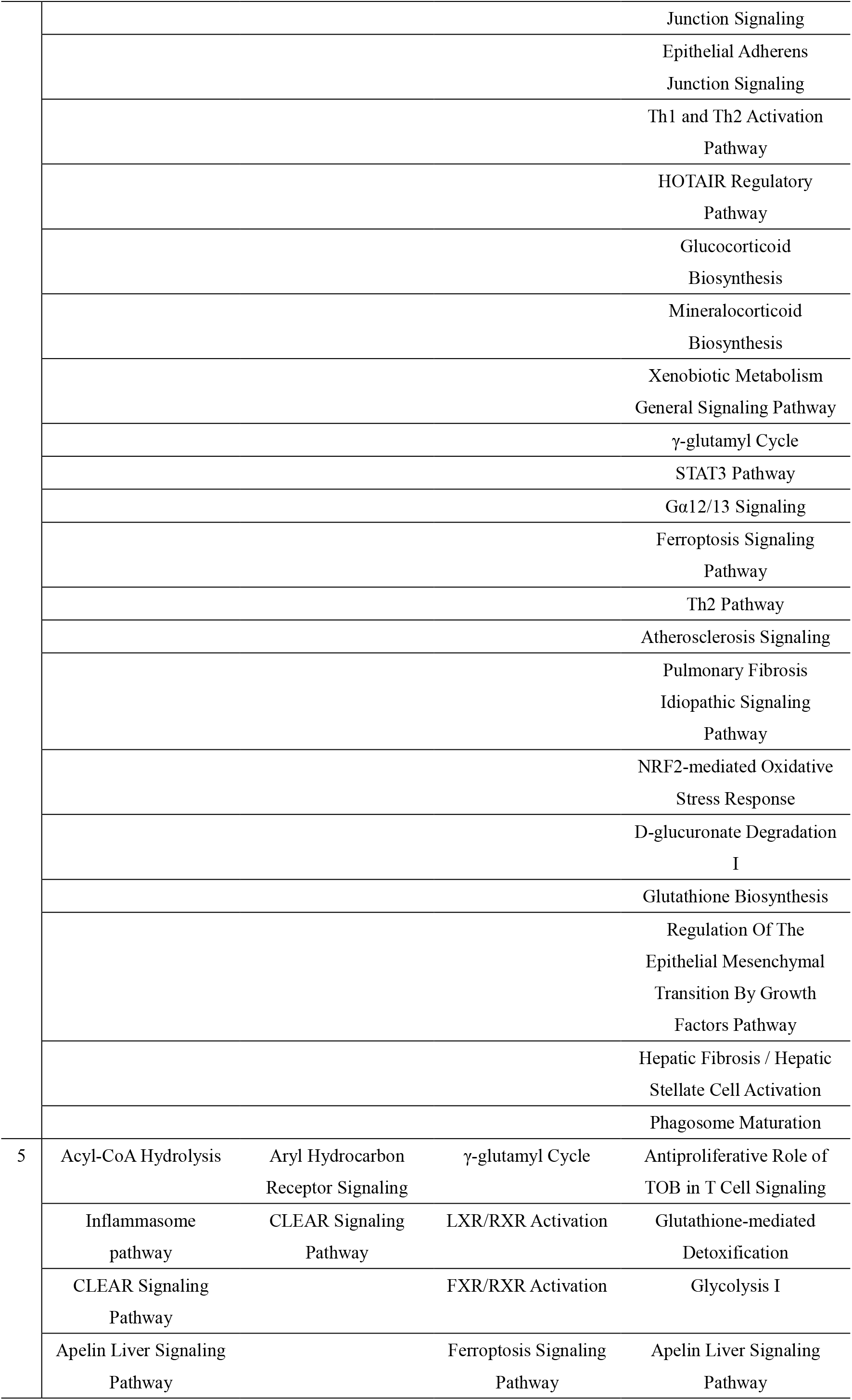
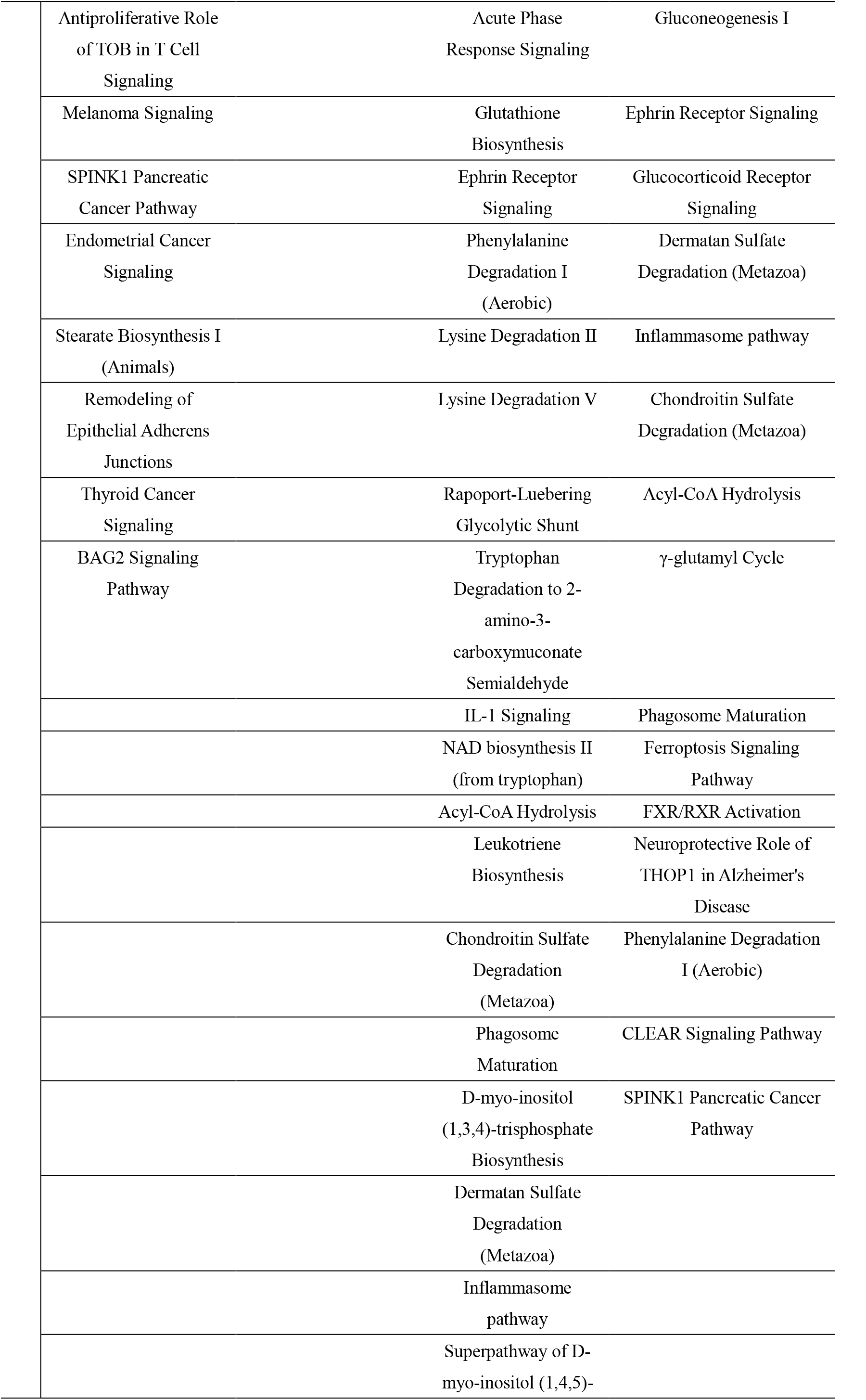
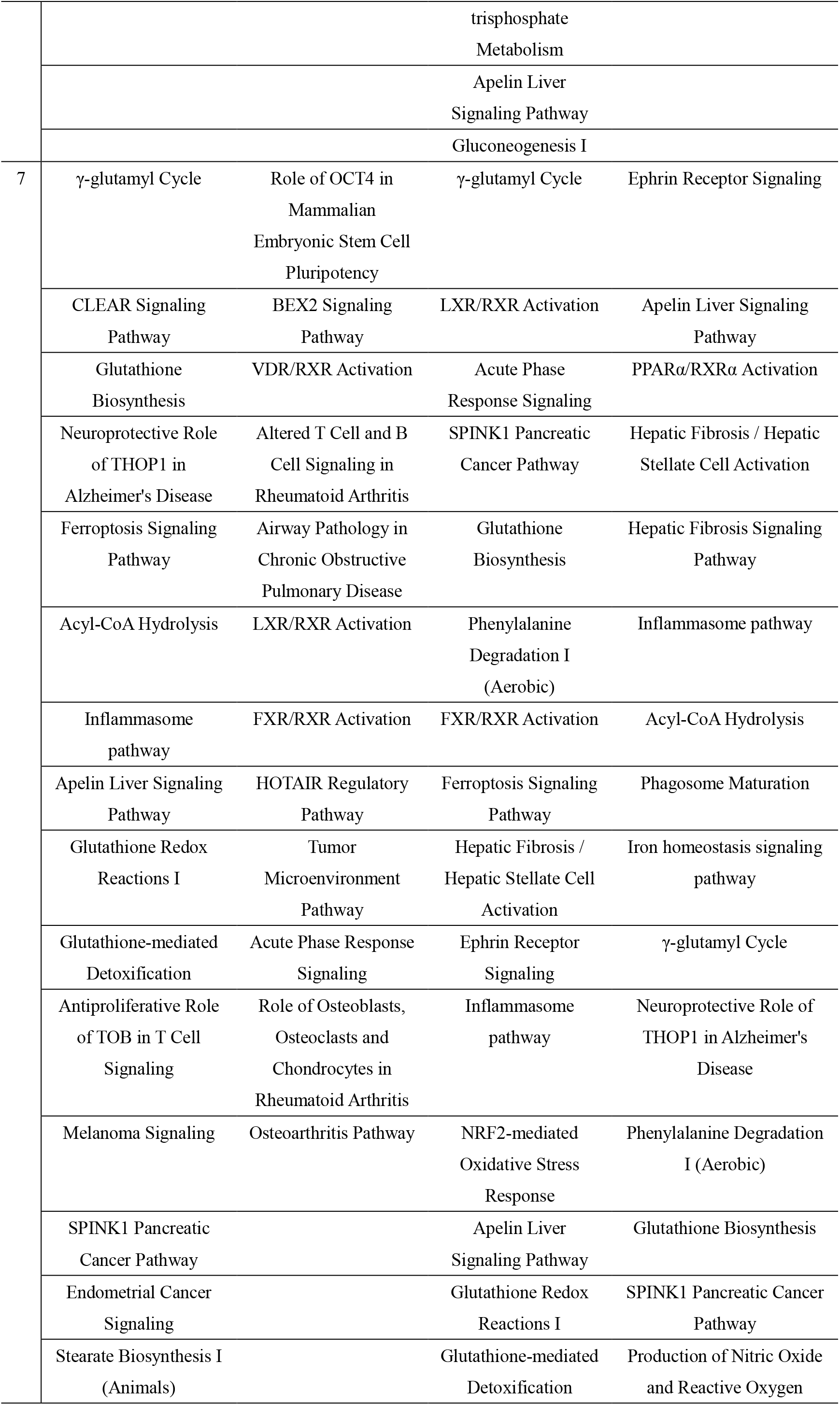
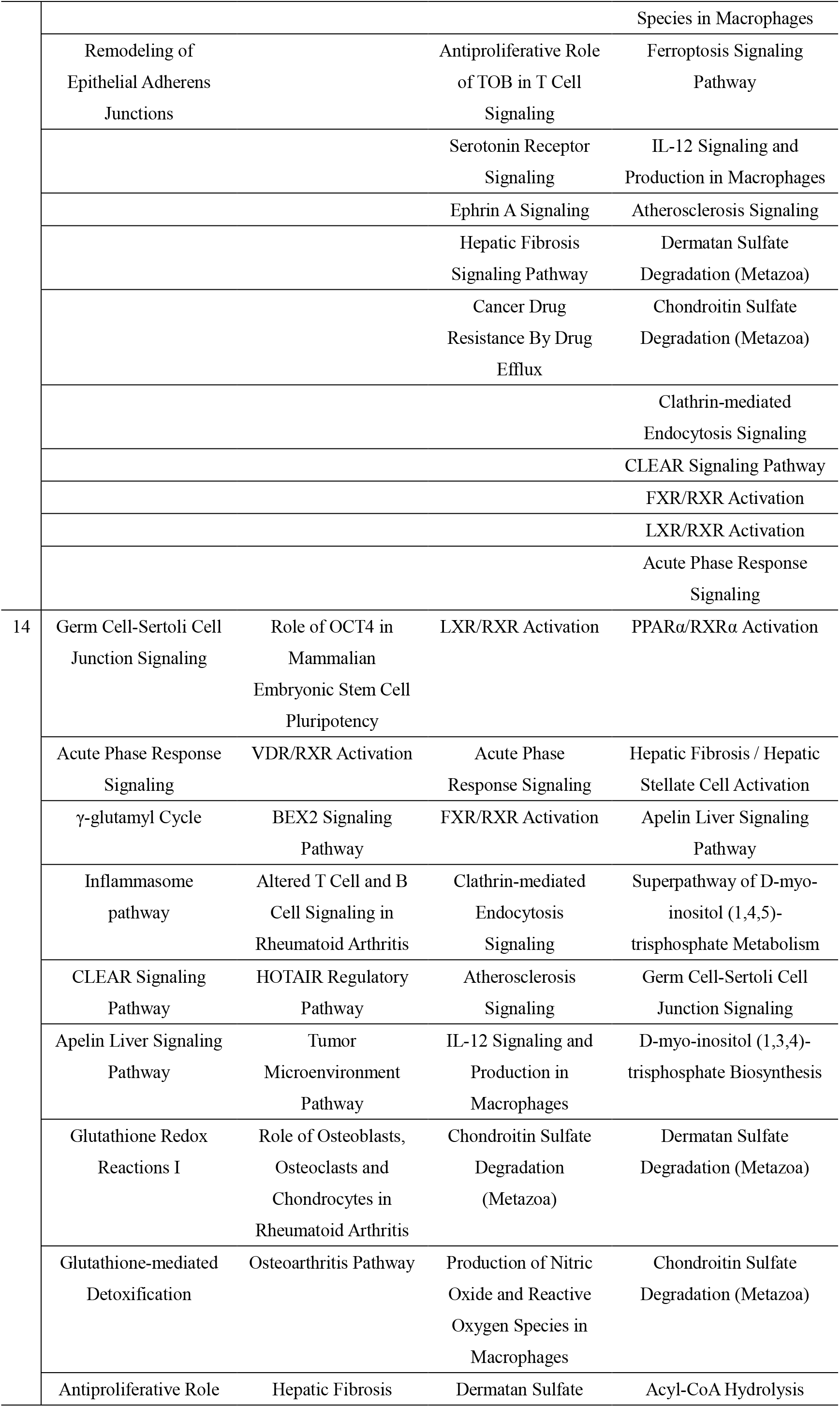
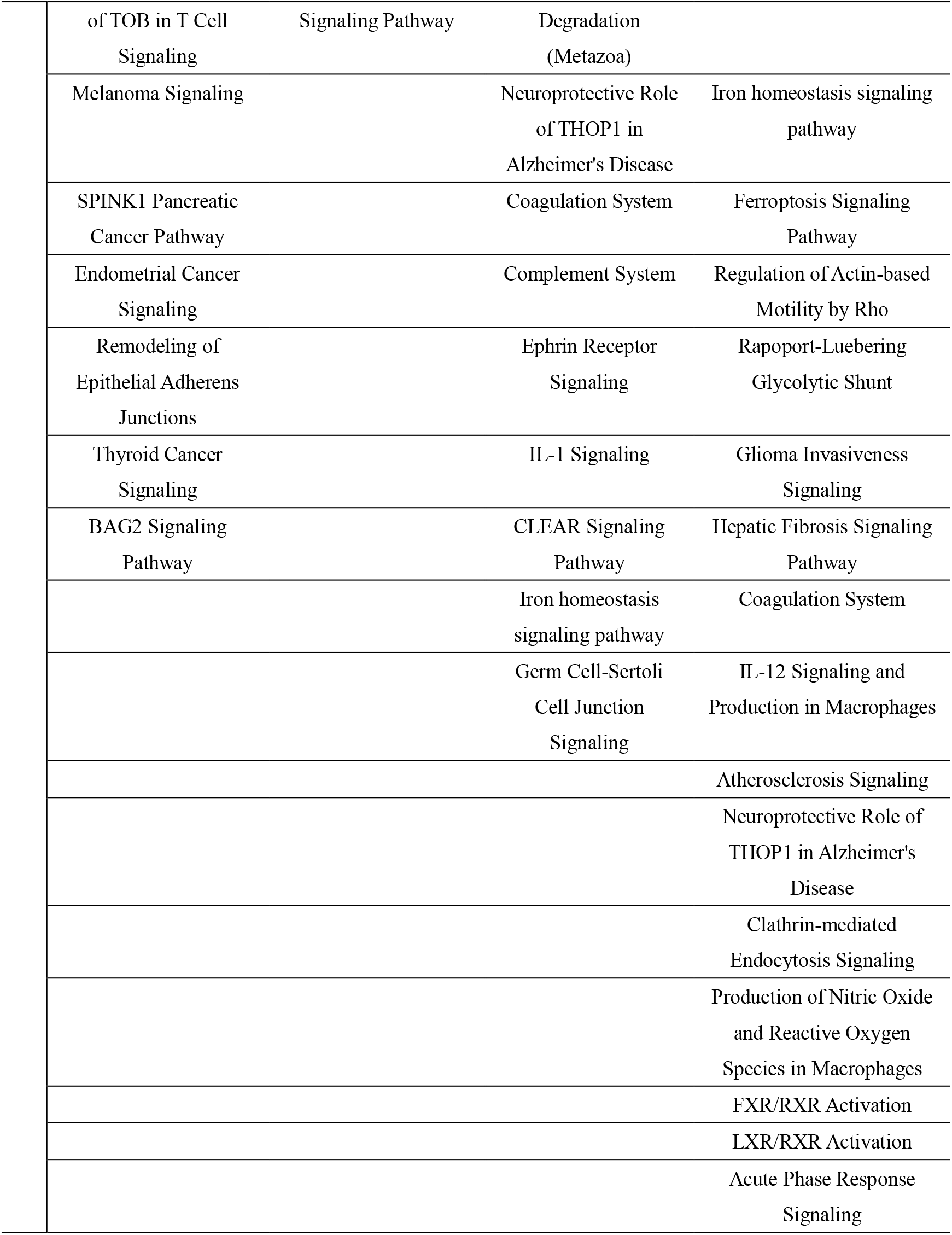
IPA pathway of rats in different groups

To intuitively see the effect of adjuvants on the immune system, we first observed the results of the comparison between the antigen–adjuvant group and the antigen– adjuvant group to explore the difference between the antigen-adjuvant group and the simple antigen group after the addition of adjuvant. As shown in Table 10, on the first day after injection, among urinary proteins, acute phase response signaling, airway pathology in chronic obstructive pulmonary disease, IL-12 signaling and production in macrophages, ferroptosis signaling pathway, granulocyte adhesion, and diapedesis, production of nitric oxide and reactive oxygen species in macrophages, and glutathione biosynthesis and inflammasome pathway were also known as immune system-related pathways. On the third day, the ferroptosis signaling pathway, acute phase response signaling, glutathione biosynthesis, airway pathology in chronic obstructive pulmonary disease, phagosome maturation, and the inflammasome pathway, which are related to the immune system and inflammation, still existed. However, in our adjuvant and antigen groups, there were no obvious immune system relationships or inflammatory pathways on Days 1 and 3. By Day 5, we observed the emergence of IL-1 signaling, which is closely associated with stimulating APC and T-cell activation, promoting B-cell proliferation and antibody secretion. At this time point, the antigenic group still did not have obvious immune system-related pathways, and the adjuvant group began to exhibit signs of the involvement of an inflammasome pathway related to the inflammatory response. After 7 days, acute phase response signaling, glutathione biosynthesis, the ferroptosis signaling pathway, and the inflammasome began to appear in the adjuvant group, and the antigen group pathway was associated with inflammation and the immune system. In the last 14 days, IL-1 signaling, IL-12 signaling, and production in macrophages were observed in the comparison between the antigen plus adjuvant and antigen groups. In the adjuvant group, acute phase response signaling and the inflammasome pathway were still involved, which may be related to the fact that the adjuvant group could not stimulate specific immunity to produce antibodies. Alterations in T-cell and B-cell signaling in rheumatoid arthritis were found in the antigen group. Through the comparison of the three groups, we noted that after performing the adjuvant–antigen plus adjuvant and antigen group comparison, protein changes related to the immune system and inflammatory reaction pathways were observed after one day, and after 5 to 14 days, antigen-presenting cells, T cells, and B-cell-related pathways appeared. Inflammatory pathways did not appear in the adjuvant and antigen groups until 7 days later, and no changes in B-cell-related pathways were observed in the adjuvant group by 14 days. When the aluminum hydroxide adjuvant was added to the antigen, as observed in urine, the adjuvant helped to stimulate the immune system to respond earlier.

We also observed changes in the urinary protein antigen plus adjuvant group compared with the control group at each time point, and we found that on Day 1, IL-17a signaling in fibroblasts, differential regulation of cytokine production in vitro in epithelial cells by IL-17a and IL-17F, the ferroptosis signaling pathway, acute phase response signaling, and the inflammasome pathway were related to inflammation. On the third day, IL-8 signaling, T-cell receptor signaling, Th1 and Th2 activation pathways, the Th2 pathway, other antigen recognition-related macrophages, T-cell activation, and B-cell proliferation pathways began to emerge. Seven to 14 days later, IL-12 signaling and production in macrophages could be observed to change the pathways related to antigen-presenting cells and B cells. The results showed that we were able to observe the process by which a vaccine stimulates the immune system and the related biological pathway changes in urine.

Overall, our results showed a series of changes in the urine during the immune response. Urinary proteins have been observed from the early inflammatory response to antigen presentation and recognition by macrophages at the antigen recognition stage to the activation and differentiation stages of immune cells, activation of the Th1 and Th2 pathways, activation of T cells, and proliferation of B cells induced by related interleukins. In addition, different immune effects of different groups are also reflected in urinary proteins. Adjuvants can better stimulate the immune response process of the immune system, while the immune response process without the addition of adjuvants can be delayed or even weakened.

These results open up new ideas for future studies of the immune system. This is an experiment in the new field of urinary proteomics and will encourage more experiments on urinary proteomics and the immune system. The changes in the immune system caused by immunogenicity can be observed in the early stages of urinary proteins, which can provide some new clues and a basis for accelerating vaccine development. These results may even provide some insights into the early detection of side effects in vaccine development.

## Notes

### Competing Interest Statement

The authors have declared no competing interest.

